# 3D spheroid-microvasculature-on-a-chip for tumor-endothelium mechanobiology interplay

**DOI:** 10.1101/2022.11.19.517181

**Authors:** Yingqi Zhang, Fengtao Jiang, Yunduo Charles Zhao, Ann-Na Cho, Guocheng Fang, Charles D. Cox, Hala Zreiqat, Zu Fu Lu, Hongxu Lu, Lining Arnold Ju

## Abstract

In the final step of cancer metastasis, tumor cells become lodged in a distant capillary bed, where they can undergo extravasation and form a secondary tumor. While increasing evidence suggests blood/lymphatic flow and shear stress play a critical role in the tumor extravasation process, there is a lack of systematic and biomechanical approaches to recapitulate sophisticated 3D microtissue interactions within the controllable hydrodynamic microenvironment. Here, we report a simple-to-use 3D spheroid-microvasculature-on-a-chip (SMAC) model. Under static and controlled flow conditions, the SMAC recapitulates the biomechanical crosstalk between heterogeneous tumor spheroids and the endothelium in a high-throughput and quantitative manners. As an *in vitro* metastasis mechanobiology model, we discover 3D spheroid-induced endothelial compression and cell-cell junction degradation in the process of tumor migration and expansion. Lastly, we examine the shear stress effects on the endothelial orientation, polarization as well as the tumor spheroid expansion. Taken together, our SMAC model offers a miniaturized, cost-efficient and versatile platform for future investigation on metastasis mechanobiology, enhanced permeability and retention effect and even personalized therapeutic evaluation.

## 1. Introduction

During cancer progression, angiogenesis is a critical driver for early stage tumor growth, and metastasis in the late stage leads to more than 90% of cancer-related deaths in patients.^[1]^ The metastatic cascade involves a multi-step process including tumor cell invasion, migration, intravasation, dissemination, extravasation, and colonization at a secondary site. These processes are often found in blood or lymphatic vessels where tumor cells are commonly subjected to shear stress. Though only around 0.01% of tumor cells that have survived from circulatory system and colonized at the surrounding tissue, it is essential to model the process of extravasation since the role of this minor population determines the metastatic potential.^[2]^

Tumor cell extravasation consists of a few steps, including adhesion, modulation of the endothelial barrier, transendothelial migration, and crossing the vascular basement membrane.^[3]^ During extravasation, both tumor and endothelial cells experience mechanical forces.^[4]^ For instance, it has been shown that metastatic breast epithelial cells reduce the stiffness of the endothelium and promote epithelial cell transmigration.^[5]^ Metastatic tumor cells have shown decreased stiffness,^[5–6]^ and their nuclei soften during transendothelial migration.^[7]^ More importantly, fluid shear stress extensively affects the extravasation process. It has been reported that low fluid shear stress caused a 2-fold increase in the efficiency of breast cancer cell adhesion to endothelium and promoted colonization of tumor cells after extravasating from the vascular system.^[8]^ Other studies suggested that a certain range of fluid shear stress can inhibit tumor cell adhesion^[9]^ and high shear force can lead to a malignant phenotype of tumor cells.^[8b]^ Collectively, these studies demonstrate that tumor metastasis is tightly regulated and coordinated by hydrodynamic forces, and the underlying mechanobiology interplay between tumor and endothelial cells is a fansinating open question to study.

Nevertheless, recapitulating the physiological and mechanical microenvironment to study tumor extravasation and metastasis remains to be a challenge in the field. Specifically, the blood vessels where tumor invasion occurs are not accessible and feasible to investigate alongside 3D tumor tissues simultaneously. Numerous studies have used trans-well plates to co-culture the cancer and endothelial cells to examine the transmigration,^[10]^ for example the molecular-to-phenotypic features of tumors for mortality and endothelial invasion.^[11]^ However, those systems still lack shear stress modulation built into the process. Endothelized microfluidics and vessel-on-chip^[12]^ have rapidly emerged as comprehensive humanized models that incorporate sophisticated vascular anatomies and spatiotemporally regulated hydrodynamic flow.^[13]^

3D multicellular tumor spheroid is a promising model that mimics the *in vivo* solid tumor. The tumor spheroids have potential to bridge the gap between a 2D monolayer cell culture and *in vivo* studies by providing similar *in vivo* biomechanical integrities, such as spatial interactions of cells with surrounding cells and extracellular matrix^[14]^ in the tumor mcroenvironment.^[15]^ In this regard, we employed a liquid dome model to produce more than 200 tumor spheroids in less than a day as previously reported.^[16]^ Specifically, the surface tension created by this model, lead to the formation of the liquid dome of cancer cell suspension on the microwell array on the chip. The liquid height at different positions of the dome controls the number of cells that settle in the microwells, producing 3D spheroids with diverse sizes, critical for recapitulating the heterogeneity of human tumor spheroid.

In this study, we developed a spheroid-microvascular-on-a-chip (SMAC) model that integrates 3D tumor spheroids into an endothelialized microfluidic device (Figure 1A). Tumor spheroids with gradient-sizes were generated using a liquid dome method. The SMAC model enabled a thorough investigation of the interaction between various tumor sizes and the endothelial layer under a diverse range of shear stresses, including; 1) tumor spheroid-induced endothelial cell compression and cell-cell junction degradation (*i.e*., VE-cadherin); 2) tumor spheroid migration and expansion in the endothelialized environment; and 3) the shear stress effects on the endothelial cell orientation and polarization as well as tumor spheroid expansion (Figure 1A-B). By seeding tumor spheroids in an endothelialized microfluidic channel, our SMAC model resembles the tumor microenvironment, in particular, incorporating an endothelial layer and fluid shear stress that regulates tumor expansion and extravasation.

**Figure 1.**
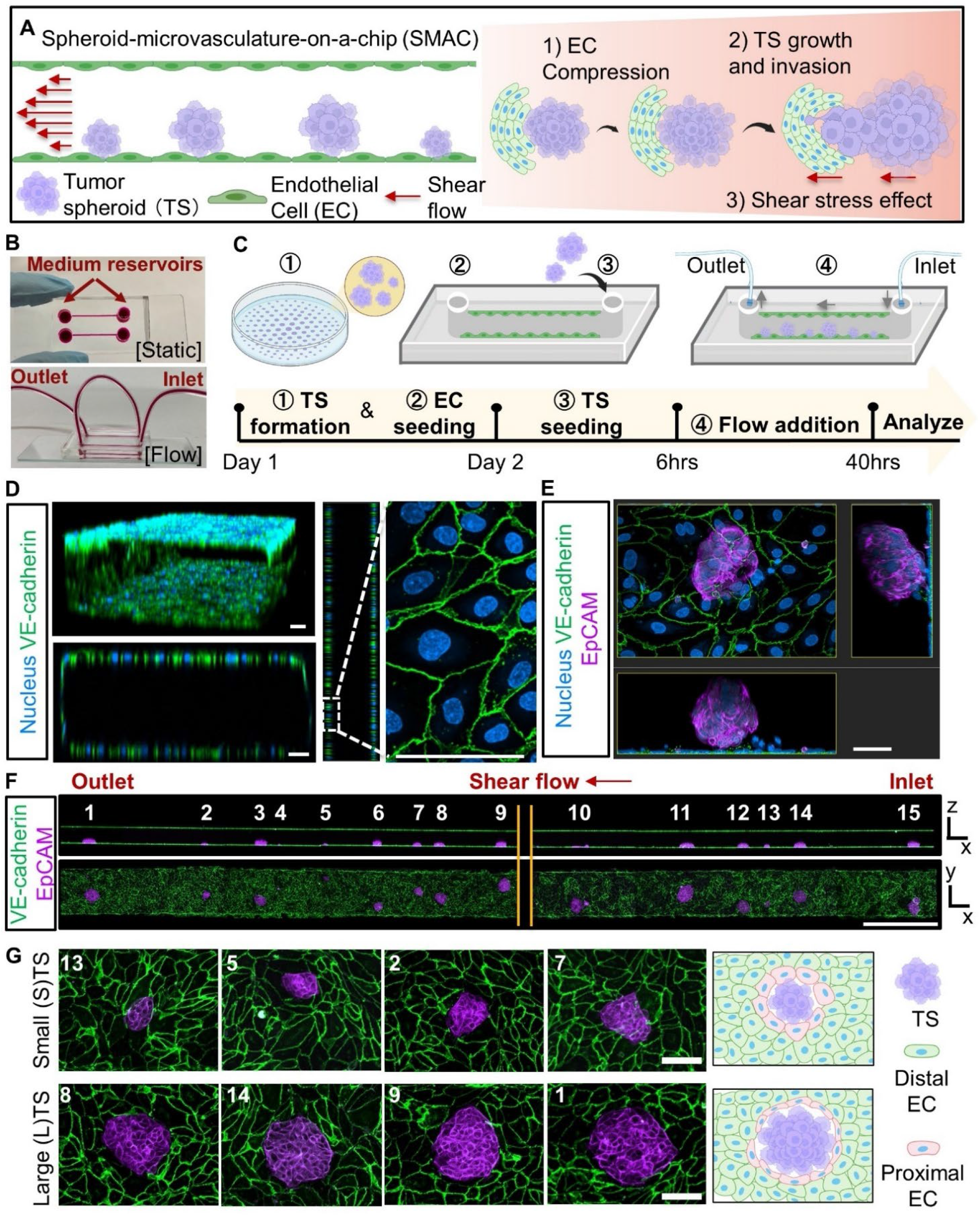
Spheroid-microvasculature-on-a-chip (SMAC) concepts and fabrication procedures. (A) Schematic illustration the SMAC to investigate tumor-endothelium mechanobiology interplay under the fluid shear stress. (B) SMAC microfluidics under static and fluid flow conditions, including medium reservoirs, inlet, and outlet. (C) Schematic workflow of SMAC fabrication in four steps. (D) Representative immunostaining images of VE-cadherin which confirm tunnel-like endothelial confluency and viability. Scale bar = 50μm by default. (E) Representative images of tumor spheroids (stained by EpCAM, the epithelial cell marker in *purple*) on the endothelial monolayer (stained by VE-cadherin in *green*). (F) The side view (top panel) and top view (bottom panel) of the full-length stitched SMAC image containing multiple tumor spheroids. Scale bar = 1mm (G) Zoom-in sperhoid images of (F). *Top*: Small tumor spheroids (STS: *D* = 30-70μm in diameter); Bottom: Large tumor spheroids (LTS: *D* = 70-150μm in diameter). Scale bar = 100μm.

## 2. Results

### 2.1. Fabrication of spheroid-microvasculature-on-a-chip (SMAC)

Using a standard soft lithography and single-mask microfabrication process,^[17]^ we created parallel microfluidic channels to model venous capillary anatomies within a few hundred microns on a PDMS chip (Figure 1B). To enable SMAC co-culture of endothelial cells and multiple tumor spheroids, we trialed multiple microchannel dimensions ranging from 300 × 100μm to 1000 × 500μm (cross setion: width × height), and then optimized the design by checking the stability and reproducibility of tumor spheroids culture in the endothelialized channel after perfusion at bulk shear stress τ_0_ = 1 to 15 dyn/cm^2^. In quick conclusion, the optimal micro-channel design was set at 600μm × 300μm × 25mm (width × height × length) with 6mm and 2mm holes punched at the inlet and outlet for static and flow culture respectively (Figure 1B, Supplementary Figure 1). This design allowed a) initial tumor spheroid localization and stable attachment in shear flow condition; b) sufficient spaces for tumor spheroid migration and expansion during the long-term culture; c) capability to culture numerous tumor spheroids in the same channel for high-throughput investigation (Figure 1B, Supplementary Figure 1).

Upon optimizing the microfluidic design, the SMAC biofunctionalization consists four major steps (Figure 1C): 1) Generate tumor spheroids using the liquid dome arrays; 2) Seed endothelial cells and form monolayers inside the microfluidic channel; 3) Seed 3D spheroids into the microvasculature-on-a-chip; and 4) Apply shear stress under perfusion. In the Step 1, human breast cancer cells, MCF-7, were seeded into agarose microwells that allow gradient-sized spheroids formation as previously reported (Figure 1C; Supplementary Figure 2A).^[16]^ After overnight culture, MCF-7 formed tumor spheroids and were harvested from microwells (Supplementary Figure 2A). Immunostaining of F-actin and nucleus by 3D confocal microscopy validated the compact intercellular connection and spherical morphology (Supplementary Figure 2B). Meanwhile for the Step 2, human umbilical vein endothelial cells (HUVEC; abbreviated as ECs) were seeded into the channels and successfully transformed into confluent monolayers spanning the entire inner lumen within 24 hours (Figure 1D). The VE-cadherin expression throughout the entire inner surface in our endothelialized microchannels further confirmed the existence of integral junctions, therefore the endothelial cells grossly achieved confluence and function appropriately (Figure 1D; *green*).^[13b]^

In the Step 3, the collected 3D tumor spheroids were then infused into an endothelialized microchannel and stabilized for 6 hours to allow firm attachment (Figure 1E). Approximately 10 to 15 tumor spheroids of different sizes (Diameter range *D* = 30 – 150μm) settled and adhered, which enabled investigation of heterogenous tumor spheroids’ interaction with endothelium (Figure 1F). To evaluate the size effects of tumor spheroids (Figure 1G), we further categorized them into groups of Small Tumor Spheroid (STS; 30μm ≤ D ≤ 70μm) and Large Tumor Spheroids (LTS; 70μm ≤ D ≤ 150μm). For shear application in the Step 4, culture medium was perfused through the entire microchannel using a peristaltic pump at a controlled bulk shear stress (i.e., τ_0_ = 5 and 15 dyn/cm^2^; cf Figures 4 and 5).

### 2.2. Tumor spheroids compress endothelium and degrade cell-cell junction upon extravastion

To examine the biomechanical impacts of tumor extravasation against the endothelium, we co-cultured endothelial cells and tumor spheroids statically in the SMAC model and fixed them after 6 hours. We defined two regions of interest: 1) The proximal inner layer of endothelium that are in immediate contact (less than two-cell body length) with the central spheroid (Figure 2A, *broken marquee*); 2) The distal endothelial region ouside the proximal perimeter and not directly contacting the spheroids (Figure 2A, *solid marquee*). Interestingly, the proximal endothelium contacting both large (LTS; Figure 2B, ①) and small (STS; Figure 2B, ②) spheroids partially lost the VE-cadherin, while the distal regions in both cases (Figure 2B, ③ and ④) displayed little difference from those intact endothelial cells in the absence of spheroids (Figure 2B, ⑤). Moreover, we normalized the VE-cadherin intensity profile across the orthogonal line region of interest (ROI) placed at 1/3 of the cell body distance. The ROI intensity of the endothelial cells proximal to both large (Figure 2C, ①) and small (Figure 2C, ②) spheroids only displayed a single peak in confirming the loss of VE-cadherin, whereas those distal to spheroids or in the absence of any spheroids displayed double peaks demonstrating the intact VE-cadherin expression in both edges (Figure 2C, ③④⑤). These analyses confirmed that the spheroid contact edges of proximal endothelial cells no longer expressed VE-cadherin, whereas the VE-cadherin expression of distal endothelial cells remained intact. In another word, tumor spheroids degraded the endothelial junctions upon its invasion.^[18]^

**Figure 2.**
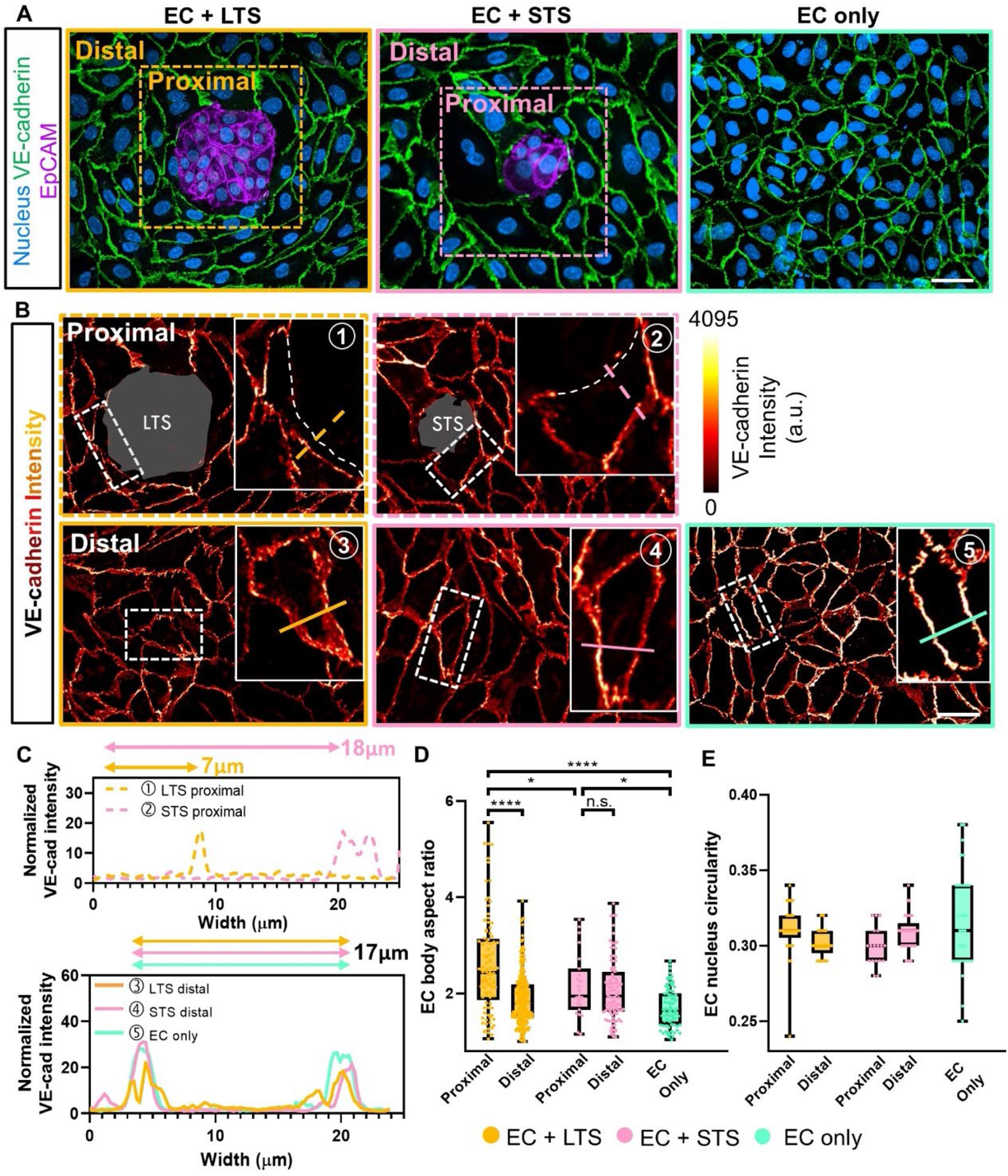
SMAC models the mechanical impacts of tumor spheroid extravastion on the endothelium. (A) Representative confocal images of endothelial cells in the prescence of a large (LTS, *left*), a small (STS, *middle*),or without (EC only, *right*) any tumor spheroid on a chip. Endothelium in the presence of tumor spheroids were classified into two subgroups – proximal and distal, which are outlined in yellow and pink marquees, respectively. Scale bar = 50μm. (B) The heat map shows VE-cadherin expression intensity of the endothelium. Scale bar = 50μm. Inset: the zoom-in images of representative proximal or distal endothelial cells in the LTS, STS or EC only settings. The white dash-line delineated the VE-cadherin signal loss. Intensity profiles in (C) were determined by the solid lines (25μm) shown in the inset. The color bar on the right corresponds to the absolute intensity level expressed in an arbitrary unit. (C) The VE-cadherin expression intensity profiles of individual ECs indicated in (B). The intensity is normalized by the minimum intensity value. (D-E) Cell body aspect ratio (D) and nucleus circularity (E) measurements. The histogram depicts all data points of n > 20 independent experiments performed in duplicate. The error bars indicate the maximum and minimum data. ns = not significant; *p < 0.05; **p < 0.01; ***p < 0.001; ****p < 0.0001, assessed by a one-way ANOVA test and Tukey’s multiple comparison test.

Intrigueingly, the proximal endothelial cells to a large spheroid were significantly compressed and elongated (Figure 2B, ①) in comparison to those proximal to a small spheroid (Figure 2B, ②). The large spheroid could compress the width of endothelial cell from 17 to 7μm while the small spheroid could not (Figure 2C, ① vs. ②⑤). However, the endothelial cells distal to both spheroid subgroups showed no compression (Figure 2C, ③ and ④)and retained their hexagonal shape comparable to those seeded without any spheroids (EC only; Figure 2C, ⑤). To quantify the compression effects, we measured the aspect ratio (AR) of endothelial cells defined by the ratio of the long axis to the short axis of the cell body. No significant difference was observed upon the aspect ratios of distal endothelial cells to large spheroids (Figure 3D, AR = 1.97 ± 0.06), small spheroids (AR = 2.07 ± 0.06) and those without spheroids (AR = 1.71 ± 0.05). However, the proximal endothelial cells to large spheroids had an increased mean aspect ratio at 2.64 ± 0.13, which decreased to 2.15 ± 0.14 for proximal endothelial cells contacting small spheroids (Figure 2D). In contrast to aspect ratio measurement, the nucleus circularity showed no significant difference among all conditions (Figure 2E). The results suggested extravasading tumor impose compressive force on adjecnt endothelial cells, and the subsequent morphological impacts depend on the spheroid sizes.

**Figure 3.**
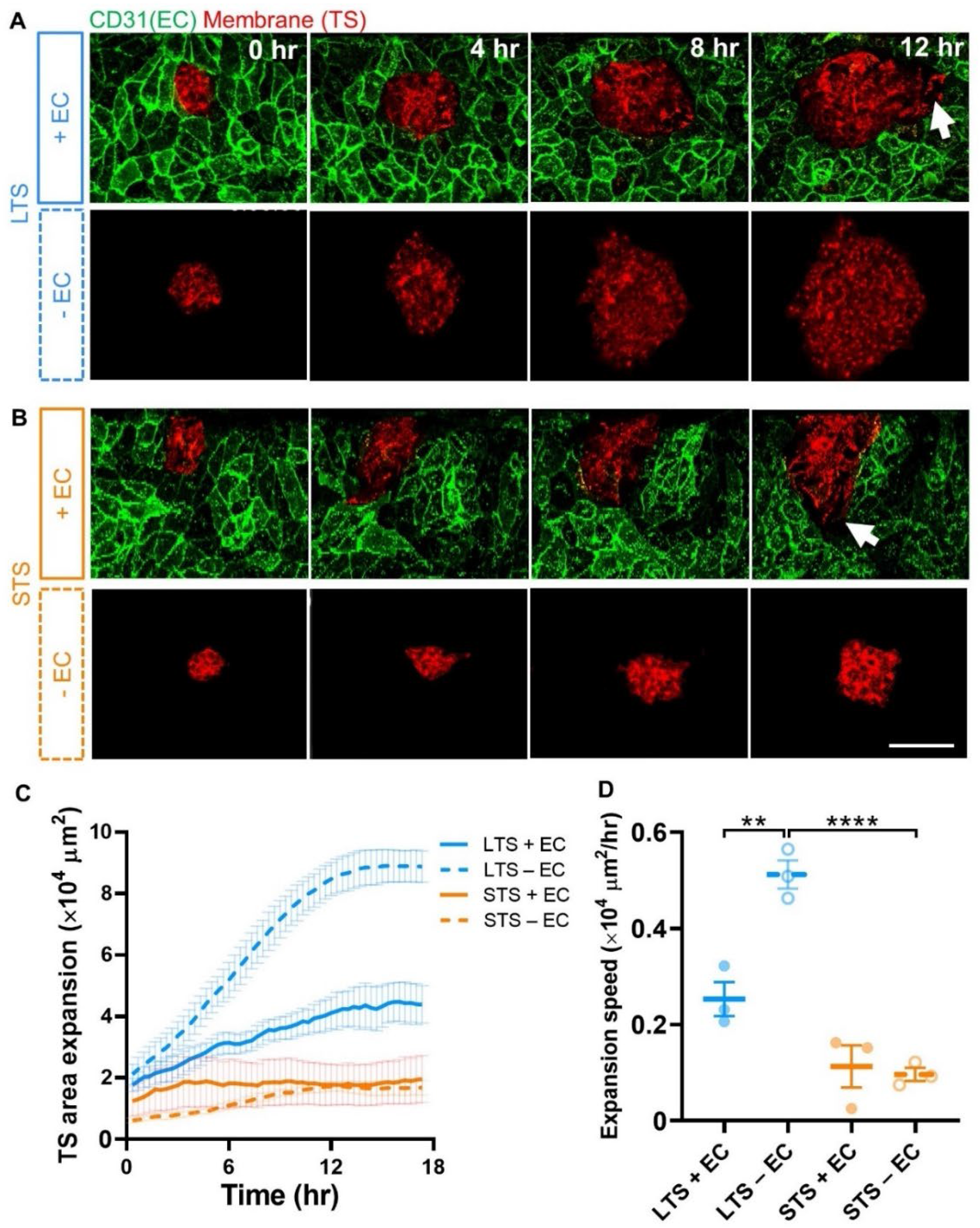
Migration and expansion of tumor spheroids are regulated by the endothelium. (A-B) Live confocal imaging of large (A) vs. small (B) spheroid in the presence and absence the endothelium. ECs were immunostained using an anti-CD31 antibody (*green*), and the cell membrane of spheroids was stained by CellMask Orange (*red*). Scale bar =100μm. (C) The area of LTS and STS spreading with or without the presence of ECs for up to 18 hours. Data shows Mean ± s.e.m. and n ⩾ 3 spheroids were evaluated in duplicate. (D) The spreading speed of LTS and STS with or without ECs over 18 hours. Data shows Mean ± s.e.m. and n ⩾ 3 spheroids were evaluated in duplicate. *p < 0.05; **p < 0.01; ***p < 0.001; ****p < 0.0001, assessed by one-way ANOVA test and Tukey’s multiple comparison test.

**Figure 4.**
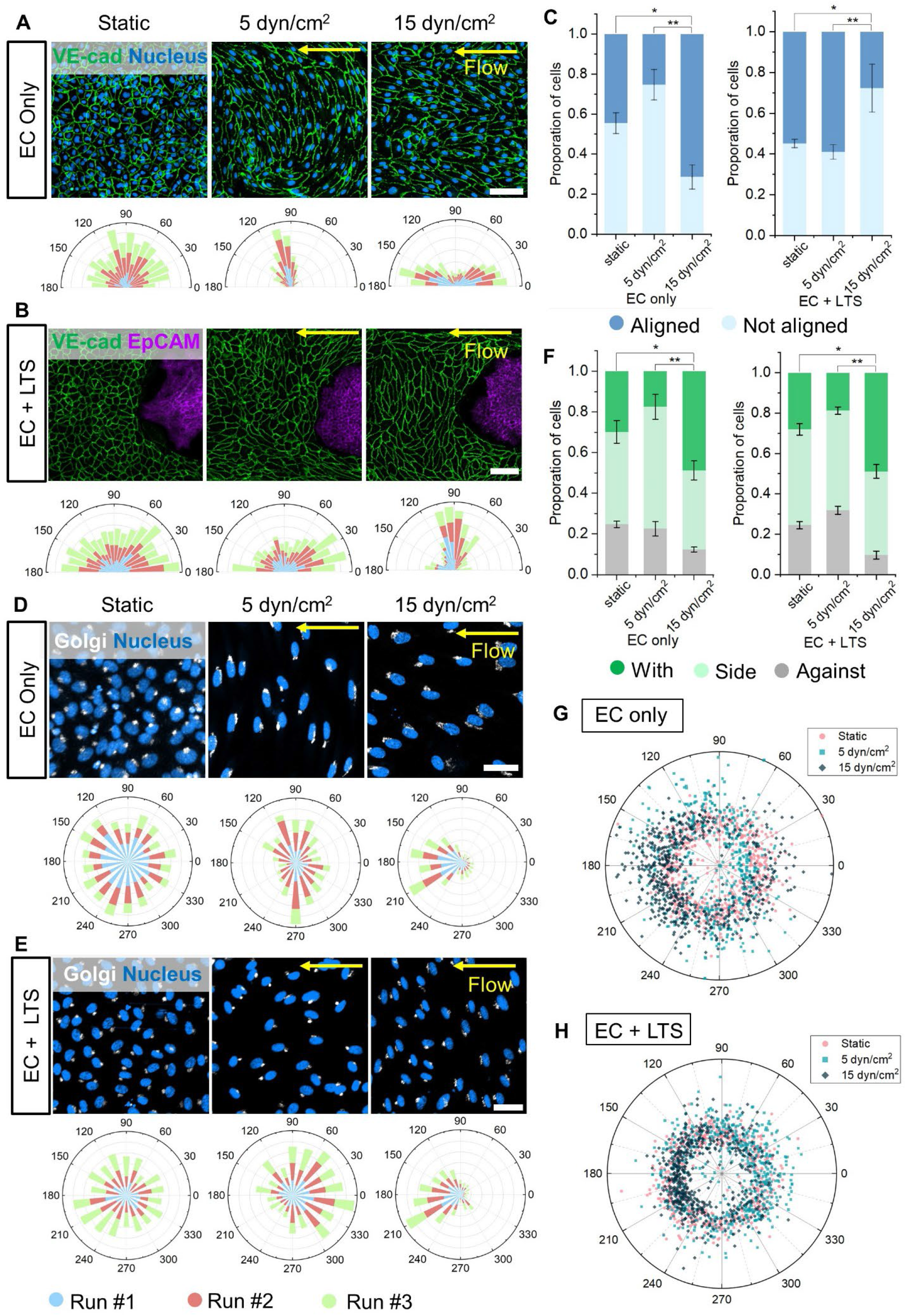
SMAC models the impact of shear stress on endothelial orientation and polarization. The endothelial orientation and polarization affected by static; 0 dyn/cm^2^ and laminar flow;5 dyn/cm^2^, 15 dyn/cm^2^. (A) Representative confocal images of EC only (*blue: nucleus; green: VE-cadherin*) when exposed to different shear stresses for 40 hours (*top*); and the corresponding EC orientation quantification in hemi-rose graphs (*bottom*) (n = 3; approximately 600-700 cells were analyzed). Scale bar = 100μm. (B) Representative confocal images of VE-cadherin (ECs) and EpCAM (tumor spheroids) exposed to shear stress for 40 hours (*top*); and the EC orientation quantification in hemi-rose graphs (n = 3; approximately 600-700 cells were analyzed). Scale bar = 100μm. (C) The propotion of ECs aligned with the flow direction (in between 45° around the flow axis, n = 3, between 600 and 700 cells were analyzed); Data shows Mean + SD. (D) Representative confocal images of EC only (*white: golgi-body; blue: nucleus*) exposed to different shear stresses for 40 hours (*top*); and EC polarization quantification in rose graphs (n = 3; approximately 600-700 cells were analyzed per replicate). Scale bar = 40μm. (E) Representative confocal images of ECs (*white: golgi-body; blue: nucleus*) near LTS when exposed to different shear stresses for 40 hours (*top*); and EC polarization quantification in rose graphs (n = 3; approximately 600-700 cells analyzed). Scale bars = 40μm. (F) Quantification of ECs polarization relative to the flow direction (n = 3; with: in between 135 and 225° around the flow axis, side: 45–135° and 225–315° against 0–45° and 315–360°; approximately 600–700 cells were analyzed); Data shows Mean + SD. *p < 0.05; **p < 0.01, assessed by a one-way ANOVA and Tukey’s multiple comparison test. (G-H) Scatter plot of the polarization of individual ECs exposed to shear stress for 40 hours without (G) and with (H) the presence of LTS.

**Figure 5.**
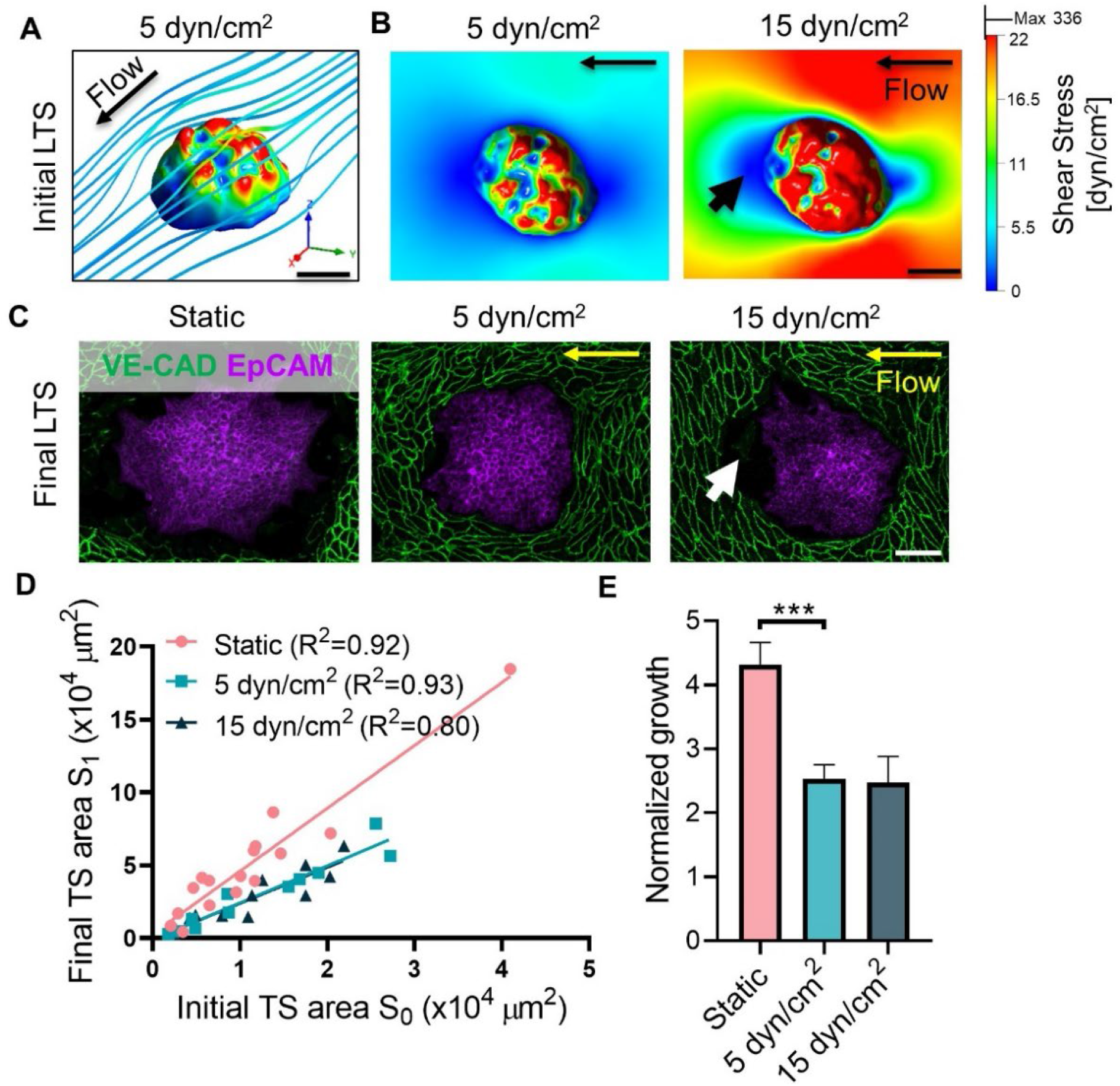
SMAC models the impact of shear stress on tumor spheroid growth. (A) 3D computational fluid dynamic (CFD) simulation showing the shear stress distribution and flow line around LTS. Scale bar = 50μm. (B) The top view of CFD simulated shear stress distribution of SMAC with LTS before adding flow. Scale bar = 50μm. (C) Representative confocal images showing EpCAM expression of LTS after 40 hours of static (left) and dynamic flow culture at 5 dyn/cm^2^ (middle) and 15 dyn/cm^2^ (right). Scale bar = 100μm. (D) Scatter graph of the final TS area vs. the initial TS area. R^2^ value is 0.92, 0.93 and 0.8 respectively. (E) The normalized growth of the TS (the slope of the line of best fit in (C)). Data shows Mean ± SD and n ⩾ 10 spheroids were evaluated in duplicate.

### 2.3. Migration and expansion of tumor spheroids are regulated by the endothelium

To track the tumor spheroid migration and invasion against the endothelium, we performed live-cell and time-lapse confocal imaging in the SMAC model (Figure 3). The endothelial cells were immunostained with the platelet endothelial cell adhesion molecule (PECAM-1 or CD31) antibody, and the tumor spheroids were labeled using CellMask Orange, referred to here as the membrane. Live-cell images were taken every 20 minutes for 18 hours (Figure 3A-B). Interestingly, at *t* = 12 hours, both large and small tumor spheroids formed branches, which migrated and extended towards the endothelial cells from the spheroids’ main bodies (Figure 2A and B, *white arrows*). In contrast, both large and small spheroids retained their spherical morphologies and exhibited isotropic expansion in the bare microchannel without endothelization (Figures 2A and B). We then quantified the area expansion of the tumor spheroids and the result showed that large spheroids with endothelium (0.25 ± 0.06 ×10^4^ μm^2^/hr) exhibited significantly lower expansion rates compared to those without endothelium (0.51 ± 0.05 ×10^4^ μm^2^/hr); while small spheroids had similar expansion rates with (0.11 ± 0.08 ×10^4^ μm^2^/hr) and without (0.096 ± 0.02 ×10^4^ μm^2^/hr) endothelium (Figure 2C-D). The results suggested that the endothelium significantly decreased the expansion rate of large but not small tumor spheroids. Moreover, the expansion rate of large spheroids was significantly higher than small spheroids without the endothelium, which was not evident in the presence of endothelium (Figure 2D). Nonetheless, the large spheroids with and without endothelium continued expanding until 12.6 and 13.6 hours, respectively; while the expansions of small spheroids with and without endothelium saturated in approximately 3.3 and 11 hours, respectively (Figure 3C). This result suggested that endothelium accelerated the saturation of tumor spheroid expansion, which was more pronounced in small than in large spheroids.

### 2.4. Shear stress influences the tumor–endothelialium interplay

To evaluate the impact of hydrodynamic forces on endothelium orientation and polarization with the presence of tumor spheroids, we subjected the SMAC to different levels of bulk shear stresses: static (τ_0_ = 0 dyn/cm^2^), venous (τ_0_ = 5 dyn/cm^2^) and arterial (τ_0_ =15 dyn/cm^2^) conditions^[19]^ for 40 hours using a peristaltic pump (Figure 4). The orientation was defined by the angle between the flow direction and the longest axis of an endothelial cell (Figure 4). In the absence of any tumor spheroids, the endothelial orientations were randomly distributed under the static condition (Figure 4A, *left*). When exposed to τ_0_ = 5 and 15 dyn/cm^2^ shear stress, the endothelial cells start aligning perpendicularly and parallelly to the flow direction respectively (Figure 4A, *middle* vs. *right*). Compared to the static condition, the proportion of parallelly aligned endothelial cells increased significantly to 71.4 ± 6.1% at 15 dyn/cm^2^ but decreased to 25.3 ± 7.6% at 5 dyn/cm^2^ due to the perpendicular alignment (Figure 4C). Further, we then analyzed the orientation of endothelial cells (~200 cells) adjacent to tumor spheroids. Endothelial cells around spheroids orientated randomly at static condition (Figure 4B, *left*; Supplementary Figure 3A, *left*). We found that the existance of spheroids increased the proportional of aligned endothelial cells to 58.9 ± 3.6% at the shear stress of 5 dyn/cm^2^ (Figure 4B, *middle*, and C; Supplementary figure 3A, *middle*, and C). However, the presence of spheroid transformed the endothelial orientation from parallel to perpendicular alignment under 15 dyn/cm^2^ shear stress (Figure 4B, *right*, and C; Supplementary figure 3A, *right*, and C).

We further examined the endothelial cell polarization which depicts the angle between the shear flow vector and the Golgi-nucleus vector^[20]^ that forms by connecting the nucleus and the center of the Golgi body (see Methods). The results revealed that endothelial cells in static culture remained randomly polarized in the absence and presence of spheroids (Figure 4D-E, *left*, and G; Supplementary figure 3B, *left*). Remarkably, endothelial cells alone mostly polarized aside the flow (~59.9%, between 45–135° and 225–315°) at 5 dyn/cm^2^ and started to polarize with the flow (~48.7%, between 135–225°) at 15 dyn/cm^2^ (Figure 4D, *middle vs. right*). The presence of large spheroids increased the proportion of endothelial cells that polarized against (~31.9%) the flow at 5 dyn/cm^2^, but had limited change in endothelial polarization at 15 dyn/cm^2^ (Figure 4E, *middle vs. right*, F and H). Similar polarization results were observed in endothelium around small spheroids (Supplementary Figure 3B and D). Taken together, these data indicated that endothelial cell orientation and polarization are highly affected by shear stresses and the tumor spheroids.

Furthermore, we evaluated the shear stress effects on the tumor spheroid expansion in the SMAC model. We reconstructed the contours of tumor spheroids derived from 3D confocal images and mapped the surface shear stress τ for both large (Figure 5A and B) and small (Supplementary figure 4A and B) SMAC models by the established computational fluid dynamic (CFD) analysis.^17a, 21]^ Notably, the mean shear stress of the large spheroid exposed to a bulk shear at τ_0_ = 5 dyn/cm^2^ (τ_ave_ = 9.7 dyn/cm^2^) was significantly lower than that at τ_0_ = 15 dyn/cm^2^ (τ_ave_ = 31.3 dyn/cm^2^) (Figure 5B, *left vs. right*). Interestingly, we found a large cavity between the endothelium and the tumor spheroids 40 hours post static culture (Figure 5C, *left*; Supplementary figure 4C, *left*), which was substantially reduced when the flow was applied (Figure 5C, *middle and right*; Supplementary figure 4C, *middle and right*). Relative to the static condition, the high shear stress at 15 dyn/cm^2^ led to a reduced cavity (~76%) formed between the tumor spheroids and the downstream endothelium (Figure 5C; Supplementary figure 4C, *right, white arrow*), possibly due to the low shear pocket found at the back of the spheroid in the CFD simulation (Figure 5B; Supplementary figure 4B, *right, black arrow*). In comparison, a much more reduced cavity (~94%) (Figure 5C; Supplementary figure 4C, *middle*) and less obvious low shear pocket (Figure 5B; Supplementary figure 4B, *left*) were observed when the shear stress at 5 dyn/cm^2^ was introduced.

We then quantified the area expansion of each individual tumor spheroids and found that the spheroid’s final area was linearly proportional to the initial area at static (τ_0_ = 0 dyn/cm^2^), venous (τ_0_ = 5 dyn/cm^2^) and arterial (τ_0_ =15 dyn/cm^2^) conditions (Figure 5D). We then constructed the spheroid growth rate by normalizing with the initial area. Remarkably, the result showed that spheroid growth rate when exposed to low (2.53 ± 0.22) and high (2.45 ± 0.41) shear stress was significantly decreased as opposed to static condition (4.32 ± 0.34) (Figure 5D and E). This indicated the suppressive effect of the shear stress on tumor migration and expansion. In conclusion, wenous shear stress (5 dyn/cm^2^) promoted endothelial alignment and polarization perpendicular to the flow direction, while arterial shear stress (15 dyn/cm^2^) induced parallel orientation and polarization of endothelial cells with the flow. However, the existence of tumor spheroids had a more significant disruption of endothelial orientation than polarization. We also found that both venous and arterial shear exposure downregulated the tumor migration and produced smaller cavity, which implicates less endothelial permeability and slow extravasation.

## 3. Discussion

Here we describe the development and characterization of SMAC model – a new microfluidic device that recapitulates key aspects of the tumor extravasation that include size scale, an endothelial monolayer cultured throughout the entire 3D inner surface of the system, and physiologically relevant hydrodynamic parameters, which can be tightly controlled and varied. Our SMAC is ideally suited for studying tumor extravasation, which is the key step for metastasis and largely dependent on the interaction between tumor and endothelium^[22]^. More specifically, this simple-to-use humanized micorsystem also presents an experimental recapitulation of this interplay and associated mechanobiology, including endothelial compression, junction loss, polarization against shear stress and rheological microenvironment.

Of note, we screened the best design sizes of the microfluidic channel from 300 × 100μm to 1000 × 500μm (width × height). Though a large channel with 1000 × 500μm caused no significant issue of co-culture, we further reduced the size of the channel for miniaturization, saving culture medium and cell usage. However, we found that a small channel with 300 × 100μm was not able to accommodate tumor spheroids in neither static nor dynamic culture due to their inabilities to settle down on the endothelialized surface, which is caused by higher shear stress from narrow space (Supplementary Figure 5). In result, we optimized the microfluidic channel to 600μm × 300μm × 25mm (width × height × length) to allow stable attachment, sufficient spaces for tumor spheroids culture in fluid shear stress.

By simply combing the gradient-driven tumor spheroid formation (liquid dome method) and the vascularized-microfluidic method, we achieved high-throughput production and analysis of the biomechanical interplay between endothelium and heterogeneous tumor spheroids with a diameter ranging from 30 to 200 μm, simultaneously. We found that tumor spheroids degraded the neighboring endothelial cell-cell junctions. This process is distinct during tumor cell transendothelial migration, where VE-cadherin was found to play a significant role in intercellular mechanotransduction.^[23]^ Moreover, we found EC compression followed by increased endothelial cell aspect ratio through direct contact with tumor spheroids during their migration and expansion. Such biomechanical compression on endothelium was more pronounced in large than small tumor spheroids. This could reflect the higher solid stress and stiffness developed in larger tumor spheroids.^[24]^ The emerging concept of VE-cadherin degradation and endothelial cell compression is not only important in the context of tumor metastasis^[18]^ but also in leukocyte diapedesis.^[25]^

The SMAC model enables real-time monitoring of tumor spheroid expansion on the endothelium. The present study demonstrated that the endothelium decelerated spheroid expansion and induced tumor invasive branches. Combing this finding with the recent highlight that the tumor cells in the invasive branches were softer, larger, and more dynamic when compared to those at the spheroid core,^[26]^ the SMAC platform would provide a comprehensive understanding on the biomechanical coordination of the tumor extravasion through the endothelium.

By introducing shear flow into the endothelialized tumor-spheroid-on-chip, we could analyze the endothelial and tumor spheroid response to different shear stresses. Venous shear stress (5 dyn/cm^2^) promoted endothelial alignment and polarization perpendicular to the flow direction while arterial shear stress (15 dyn/cm^2^) induced parallel orientation and polarization of endothelial cells with the flow. However, the existence of large tumor spheroids had greater disruption of the endothelial orientation than the polarization.

During intravasation and extravasation, tumor cells need to develop mechanisms to resist the fluid shear forces, tether to the endothelium, and migrate across the endothelial boundary to colonize at the distant site. It is suggested that interstitial, vascular, or lymphatic fluid shear stress can promote tumor metastasis^[11b]^. Shear stress induced activation of the mechanosensitive ion channels such as Piezo1^[27]^ and clustering of mechanoreceptors such as integrins^[28]^ were highlighted in cancer metastasis. In addition to tumor cell extravasation, leukocytes extravasation are also regulated by fluid shear stress,^[25]^ which increases endothelial plasma membrane tension and activates the Piezo1.^[29]^

More intriguingly, we noticed that the cavity between an expanding tumor spheroid and the adjacent endothelium was significantly suppressed by shear stress as opposed to the static condition. Considering that the cavity reflects the leaky microvasculature induced by the tumor, this finding possesses high implication to the tumor enhanced permeability and retention (EPR) effect which is known as the principle of nanoparticle design.^[30]^ Taken together, the SMAC model offer a convenient means for studying the extravasation mechanobiology in multiple contexts and shed light to future mechanomedicine drug discovery.^[31]^

## 4. Experimental Section

### 4.1. Microfluidic chip fabrication

The microfluidic devices were designed and optimized for dimension enabling high-throughput production. After being designed in AutoCAD@ software (version 2015; AutoDesk, San Rafael, CA, USA), the microchannel master was fabricated using dry photoresists.^[17a]^ Firstly, a 6-inch silicon wafer was cleaned and baked for 10 minutes at 200°C, followed by hexamethyldisilazane vapor priming for 30 seconds at 120°C. A layer of 200-μm constant height dry photoresist film (DJ micro laminate SUEX®) was then laminated on the silicon wafer at 65°C. Then a 100-μm dry photoresist film was then laminated similarly to the previous photoresist, followed by patterning using a direct lithography writer (Heidelberg MLA-100, 4,500 mJ cm^-2^) to create a channel with constant height at 300-μm. Upon lithography completion, the wafer was baked for 5 minutes at 90°C, followed by PGMEA rinsing until excess photoresist was completely removed. To prevent permanent PDMS adhesion, the patterned wafer was treated with silane in a vacuum for 2 hours and became the master mold.

To fabricate the microfluidic device, polydimethylsiloxane (PDMS) (Sylgard® 184 by Dow Corning) was mixed with the curing agent at a 10:1 ratio (w/w). Then the PDMS was poured on the master mold and heated in the oven at 60°C for 4 hours. After it was solidified, the cured PDMS was peeled off from the silicon master and cut into pieces. Then the inlets and outlets of the PDMS chips were punched with Ø6mm and Ø2mm biopsy punchers (World Precision Instruments) for static and flow experiments, respectively. Lastly, the PDMS chip was permanently bonded to 1mm glass slides by plasma treatment for 3 minutes.

### 4.2. Endothelization of microfluidic chips

Human umbilical endothelial cells (HUVECs) were obtained from Thermo Fisher Scientific and cultured with EGM-2 medium (EGM™-2 BulletKit™, Lonza). Once reached 80-90% confluency, HUVECs (passages 3-7) were washed with phosphate-buffered saline (PBS, ThermoFisher) and detached by trypsin/EDTA solution (ThermoFisher). After centrifuge, HUVECs were resuspended in EGM-2 medium at a seeding density of ~5 × 10^6^ cells/ml. Prior to HUVECs seeding, the microfluidic chip was sterilized with 80% ethanol for 20 min and washed thrice with PBS. Then the entire channels were coated with 100 μg/mL human plasma fibronectin (Thermo Fisher), incubating in a 4°C fridge overnight. The channels were then rinsed with PBS twice, and then 5 μl of prepared HUVECs suspension was injected into each microchannel. The microfluidic chip was then immediately flipped upside down to allow HUVECs attachment to the channel’s upper surface for 20 minutes. Then the chip was flipped again to allow HUVECs to attach to the bottom surface for 20 minutes. After that, the EGM-2 medium was added to the reservoirs to culture HUVECs statically overnight, and the endothelialized microfluidic chip was completed.

### 4.3. Tumor spheroid preparation and seeding into microwells

The gradient-sized multicellular tumor spheroids were prepared using the reported method.^[16]^ Briefly, MCF-7 cells were obtained from Australia Cell Bank were cultured with RPMI1640 medium supplemented with 10% fetal bovine serum (FBS, Sigma-Aldrich) and 100 U/ml penicillin/streptomycin. When the cells were up to 70%~80% confluent, MCF-7 cells were washed with phosphate-buffered saline (PBS) and detached by trypsin/EDTA solution. After centrifuge, MCF-7 cells were resuspended in RPMI1640 medium at a seeding density of 5 × 10^6^ cells/ml. Then, 100ul of MCF-7 cell suspension was added on top of the 1.8% agarose chip (containing 217 microwells) in a 6-well plate and let settle for 5 minutes. Afterward, 2ml of RPMI1640 medium was added to the 6-well plate, and the MCF-7 was cultured in the agarose chip for 24 hours. Due to the difference in cell suspension volume, the number of MCF-7 cells in each microwell varies, with more cells in the middle and fewer cells towards the side of the chip. Lower concentration would lead to smaller spheroids. The MCF-7 spheroids are then retrieved from the agarose chip to a microtube for usage later.

### 4.4. Co-culture of tumor cells and HUVECs on the chip under static and medium flow conditions

After 24 hours, the MCF-7 spheroids formed and were harvested into a microtube. Once the spheroids were settled down at the bottom of the microtube, remove the supernatant and add 1ml of the EGM-2 medium. When the spheroids are settled again, inject 10 μl of the spheroid suspension into the endothelialized channel and allow 6 hours for the spheroids to stay firmly attached. For static culture, the culture medium was then added to the reservoirs and changed once a day. For dynamic culture, the tubings were connected with the microfluidic chip and the peristaltic pump (Harvard, P70) which was set at 375 μl/min and 1.12 ml/min for 5 dyn/cm^2^ and 15 dyn/cm^2^ respectively. The microfluidic chip was then cultured for 40 hours continuously.

### 4.5. Immunostaining and confocal microscopy

The SMAC was fixed with 4% paraformaldehyde (company), then thoroughly washed with PBS and permeabilized with 1%Triton X-100 (Roche) at room temperature for 1 hour. Then the chip was blocked with 5% bovine serum albumin (BSA, Sigma-Aldrich). Subsequently, the SMAC were incubated with anti-VE-Cadherin antibody (Invitrogen) conjugated with Alexa Fluor™ 488 (1:100) for endothelial cells and anti-EpCAM antibody (Cell Signaling Technology) conjugated with Alexa Fluor™ 555 (1:100) for MCF-7 tumor spheroids for 1 hour at 37°C. After washing with PBS, the chip was stained with Hoechst 33342 (Abcam) or Phalloidin (Abcam) for nuclei and F-actin for 20 minutes at room temperature, respectively. After washing, the microfluidic device was imaged using an Olympus FV3000RS laser scanning confocal microscope and operated by FLUOVIEW software.

### 4.6. Endothelial cell morphology quantification

#### 4.6.1. Aspect ratio analysis

To quantify the influence of the tumor spheroid on the endothelium, we calculated the aspect ratio of endothelial cell body and the circularity of the nuclei. The aspect ratio was defined as the ratio of the long axis to the short axis automatically calculated by IMARIS (Bitplane AG, 9.0.1, Oxford Instruments). The circularity of a cell was defined by the ratio of the cell surface area of a sphere (with the same volume as the given particle) to the cell surface area of the particle (Wadell 1932), which was also automatically calculated by IMARIS. The histogram graphs were plotted in Figure 2D-E.

The auto-detection of cells performed by IMARIS is in need for manual editing to have a better detection efficiency (correct detection of cell nuclei, shapes, and area, etc.). We employed two different image processing methods for 2D and 3D confocal image. For 2D images, the original images containing VE-cadherin, nuclei, and F-actin signals were first transported to the format of TIF through IMARIS, and then imported into Adobe Photoshop CC 2019. Due to the compromised image resolution, the defects of VE-cadherin signals at boundary of the endothelial cells have brought difficulty in detecting the whole cell body. Therefore, we manually edited the boundary of the cells according to the F-actin signals. To avoid false dectection by IMARIS, we manually added a fake nucleus at the empty space where endothelial cells were not present which were then filtered out by selecting the regions of interest.

#### 4.6.2. Orientation and polarization analysis

We quanfied the endothelial cell orientation by calculating the angle between the flow direction and the ‘cell vector’ determined by the long axis of each endothelial cell. The orientation of the endothelial cells (~200 cells for one sample, n=3) was quantified and divided into 18 subgroups from 0° to 180° and plotted into the hemi-rose graph as shown in Figure 5A-B. To plot the histogram graph in Figure 5C-D, the endothelial orientation was categorized into aligned (0-45° and 135-180°) and unaligned (45-135°) cells with the flow direction.

To quantify the polarization of the endothelial cells, we defined the ‘polarization angle’ as the angle between the vector of flow direction and the vector formed by the nucleus and Golgi body. The polarization of the endothelial cells (~200 cells for one sample, n=3) was quantified and divided into 18 subgroups from 0° to 360° and plotted into the wind rose graph as shown in Figure 5E-F. Similarly, to plot the histogram graph presented in Figure 5G-H, we categorized the endothelial cells into three types: aligned with (135-225°), side (45-135°, and 225-315°), and against (0-45° and 315-360°) the flow direction. The angle calculation for orientation and polarization was automatically generated from the detection of IMARIS and a handmade script in EXCEL (Microsoft Office 2016) which was validated by ImageJ (National Institutes of Health, USA, 1.53K). The graphs were plotted using Origin (OriginLab, 2019b).

### 4.7. Computational fluid dynamic analysis

The CFD simulation was performed over a virtual microfluidic channel (length *x*_0_=1000 μm, width *y*_0_ = 600 μm, and height *z*_0_ = 300 μm) utilizing ANSYS®Fluent 2020 R1 software (version 20.1; Canonsburg, PA, USA) as previously described.^[21, 32]^ The tumors were reconstructed from the 3D confocal scan using Imaris software (Bitplane)^[17a]^ and placed in the middle of the channel. The tumor and the channel were then meshed into 1 μm grids for an iterative solving process. The fluidic dynamic properties of the culture medium was set at a density of 1000 kg m^-3^ and a viscosity of 0.72 Pa s. The whole interstitial fluid was assumed laminar and incompressible, and the channel was assumed to be rigid. In the solving process, the standard second-order scheme and the second-order upwind scheme were selected with the COUPLED algorithm utilized pressure-velocity coupling.^[33]^ The bulk shear stress *σ*_0_ at the micro-channel inlet was set at 5 and 15 dyn/cm^2^. After solving the Navier-Stokes equation for the whole interstitial space, the simulated shear stress contours and fluid streamlines around the tumor were constructed and exported as the results.

### 4.8. Live cell imaging and analysis

For live cell imaging, before seeding the tumor spheroids, the endothelialized channel was stained with anti-CD31 antibody conjugated with Alexa Fluor™ 488 (AbCam, 1:100), and the tumor spheroids were stained with CellMask™ Orange (ThermoFisher, 1:1000) for half an hour at 37°C. After washing with PBS, the stained tumor spheroids were seeded into the endothelialized chip for live cell imaging using an Olympus FLUOVIEW FV3000 confocal laser scanning microscope adapted with Tokai Hit Incubator for 18 hours with 20 minutes intervals. To track the area expansion of tumor spheroid, the bottom layer of the tumor was extracted and manually selected using the polygon selection tool in ImageJ. The tumor spreading area was then measured in ImageJ using the measurement tool.

### 4.9. Statistical analysis

All graphical data are presented by GraphPad Prism 9.0. Statistical differences between each group were tested by a one-way ANOVA test. A p-value below 0.05 was accepted as significant.

## Conflicts of interest

There is no conflict to declare.

## Acknowledgements

We thank Dr Freda Passam, Dr Yogambha Ramaswamy, Dr Yao Wang and Jinyuan Vero Li for the helpful discussion. We thank Vivian Cheng, Wing-tim Choi for their support on imaging and device fabrication. We also thank Jacky He and Nadia Court for microfluidic design and assistance in soft lithography. This work was conducted (in part) using the Research Prototype Foundry core research facilities at the NSW node of the Australian National Fabrication Facility (ANFF-NSW), a company established under the National Collaborative Research Infrastructure Strategy to provide nano- and micro-fabrication facilities for Australia’s researchers.

This work was supported by the Australian Research Council (ARC) Discovery Project (DP200101970 – LJ), the National Health and Medical Research Council (NHMRC) of Australia Ideas Grant (APP2003904 – LJ), NSW Cardiovascular Capacity Building Program (Early-Mid Career Researcher Grant – LJ), Sydney Research Accelerator prize (SOAR – LJ), NSW CVRN-VCCRI Research Innovation Grant and Ramaciotti Foundations Health Investment Grant (2020HIG76 – LJ), Charles Perkins Centre Early to Mid-Career Researcher Seed Funding Grant (EMCR – YZ), Cardiovascular Initiative Catalyst Award seed funding (CVI – YZ). Lining Arnold Ju is an ARC DECRA fellow (DE190100609).

## Table of contents

ToC text:

We report a 3D spheroid-microvasculature on a chip (SMAC) model to recapitulates the mechanobiological crosstalk between heterogeneous tumor spheroids and the endothelium. We discover 3D spheroid-induced endothelial compression and cell-cell junction degradation in the process of tumor migration and expansion. Lastly, we examine the shear stress effects on endothelial orientation, polarization in the absence and presence of tumor spheroids.

ToC figure:

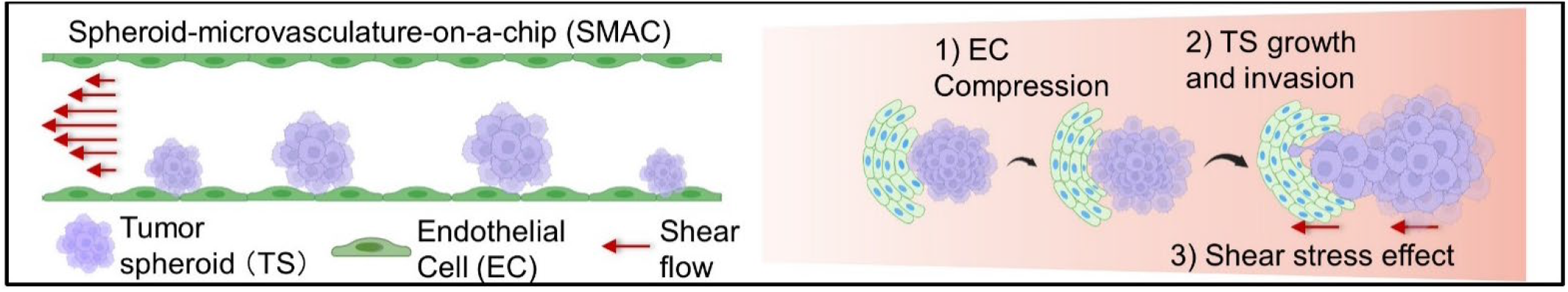

**Biographies**

**Dr Lining (Arnold) Ju**

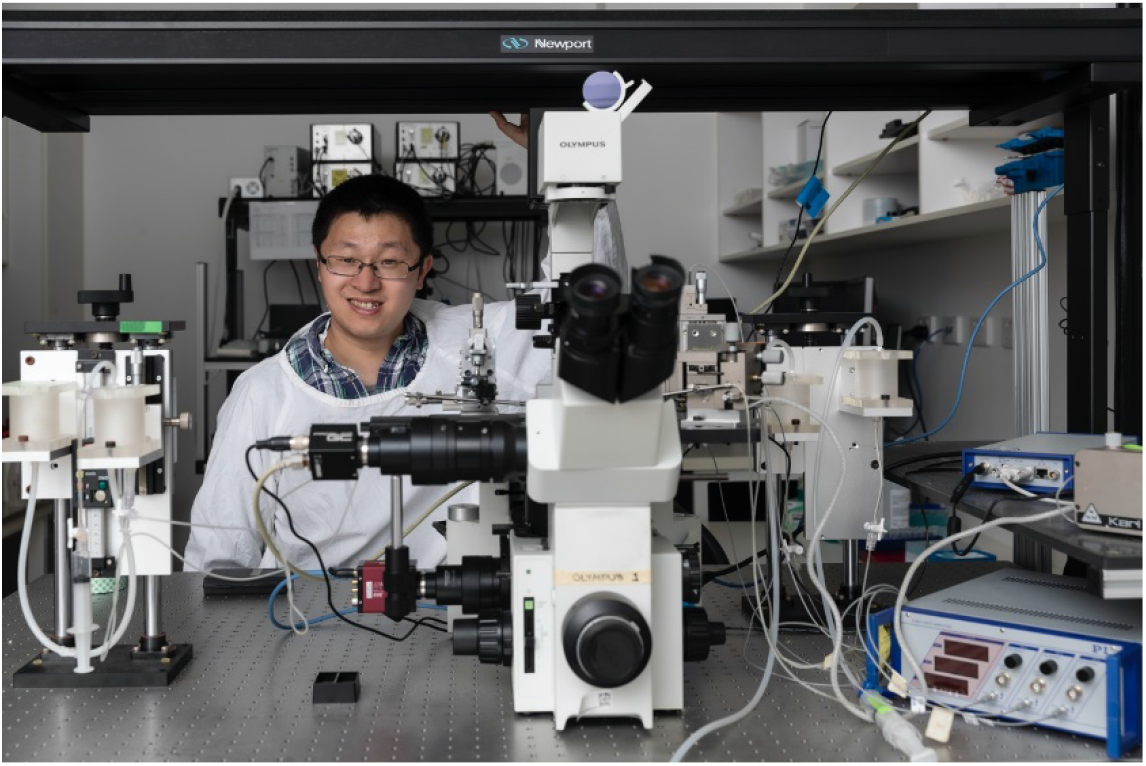

Dr Lining (Arnold) Ju received his PhD in Biomedical Engineering at Georgia Institute of Technology and Emory University, USA. In 2014, he joined the Australian Centre for Blood Diseases, Monash University, Melbourne as a junior postdoc; and relocated in 2015 to Sydney, to join the Heart Research Institute.

In 2020, Dr Ju joined the University of Sydney’s School of Biomedical Engineering as a senior lecturer and started up the Mechanobiology and Biomechanics Laboratory (MBL). By working with Charles Perkins Centre, Digital Science Institute, Royal Prince Alfred Hospital, MBL is now developing point-of-care microdevices, artificial intelligence imaging modalities and mechano-medicine screening platform to predict, diagnose and treat thrombosis.

**Dr Hongxu Lu**

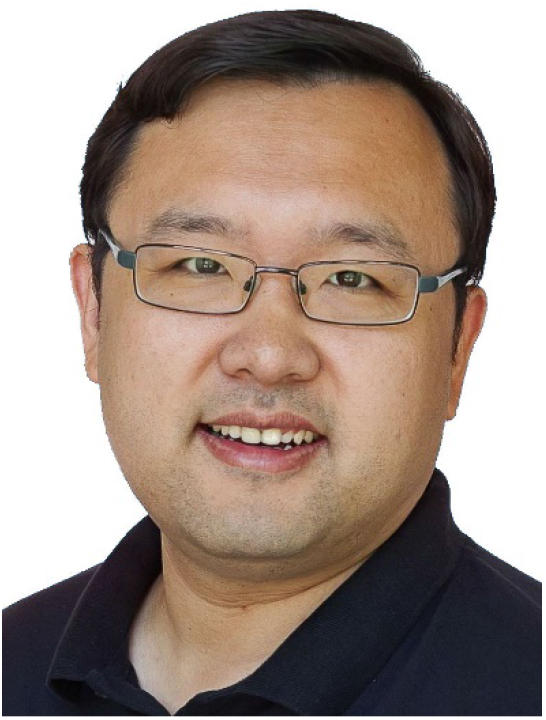

Dr Hongxu Lu obtained his PhD in 2009 from the University of Tsukuba. Then, he received postdoctoral training at the National Institute for Materials Science of Japan and the University of New South Wales (UNSW). Since 2015, Dr Lu has worked as a research fellow at UNSW, the University of Technology Sydney, and the University of Sydney. He joined the Shanghai Institute of Ceramics Chinese Academy of Science as a principal investigator in 2022. His current research focuses on developing organoid and organ-on-chips models combining bioactive materials, stem cells, 3D in vitro culture, and microfluidic technology.

**Miss Yingqi Zhang**

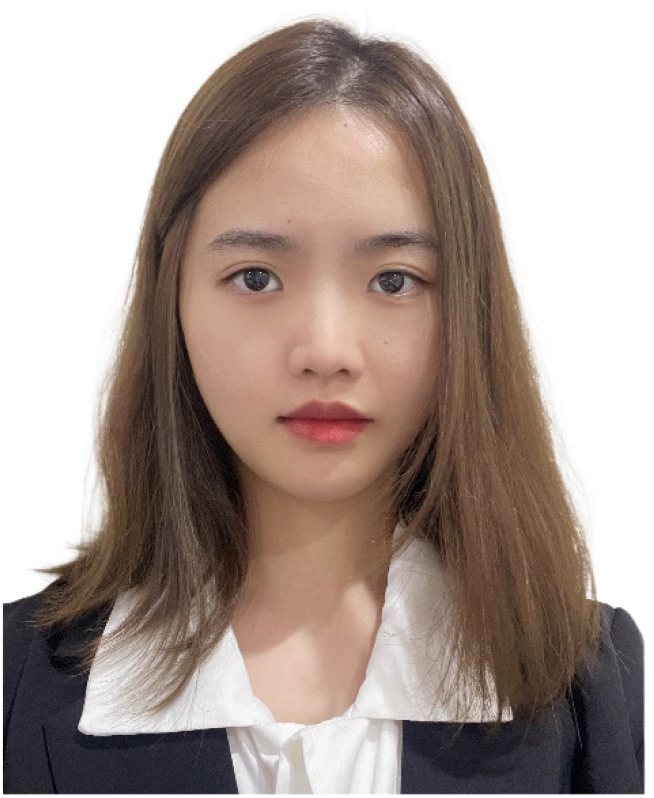

Miss Yingqi Zhang received her Bachelor of Biomedical Engineering Honour degree with first-class honour in 2020 at the University of Sydney. During the bachelor’s degree, Ms Zhang was affiliated as an intern with the Dental Research Institute at Westmead Hospital and was awarded the Faculty of Engineering Summer Research Scholarships. In 2021, Ms Zhang was awarded a full ARC PhD scholarship and started her PhD degree to develop organ-on-chip microfluidics for cancer-associated thrombosis under the supervision of Dr. Lining Arnold Ju, Dr Hongxu Lu and Dr. Yogambha Ramaswamy.

## Notes

### Competing Interest Statement

The authors have declared no competing interest.

