## Supplemental information for "3D spheroid-microvasculature-on-a-chip for tumor-endothelium mechanobiology interplay"

A

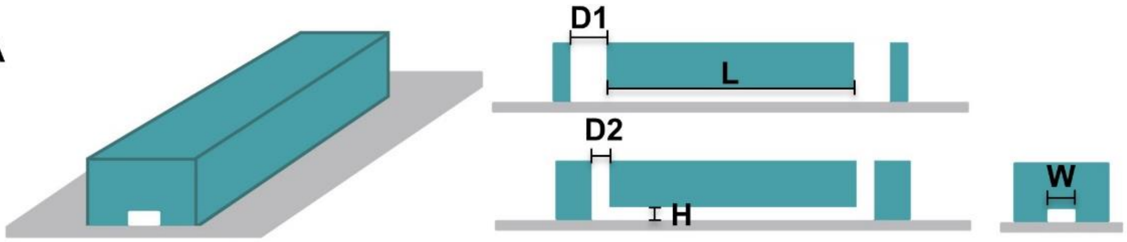


| | | |
|---|---|---|
| **L** | Total length of channel | 25 mm |
| **W** | Width of channel | 600 µm |
| **H** | Channel height | 300 µm |
| **D1** | Diameter of channel reservoir for static culture | 6 mm |
| **D2** | Diameter of channel reservoir for dynamic culture | 2mm |

B

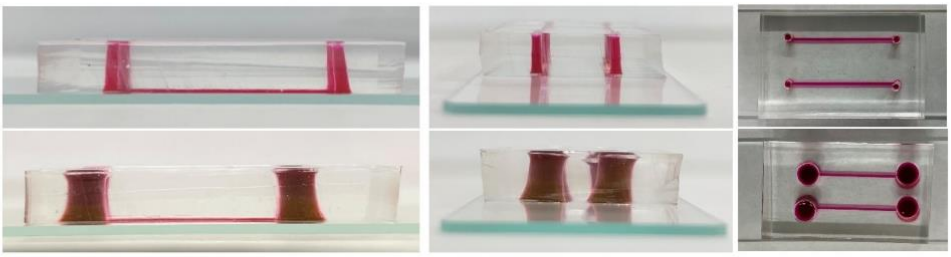

**Supplementary figure 1.** (A) Schematics of the microfluidic channel and its dimension. (B) Images of the SMAC device from different views.

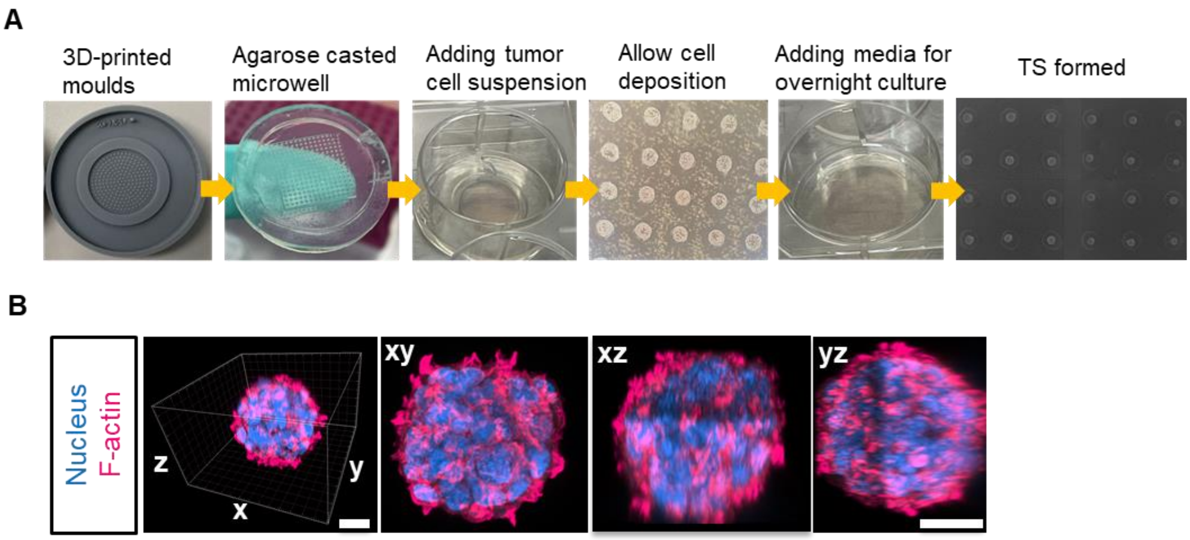


**Supplementary figure 2.** (A) Workflow for tumor spheroid formation. (B) Representative confocal images of the nucleus (*blue*) and F-actin (*magenta*) of MCF-7 spheroids. Scale bar = 30µm.

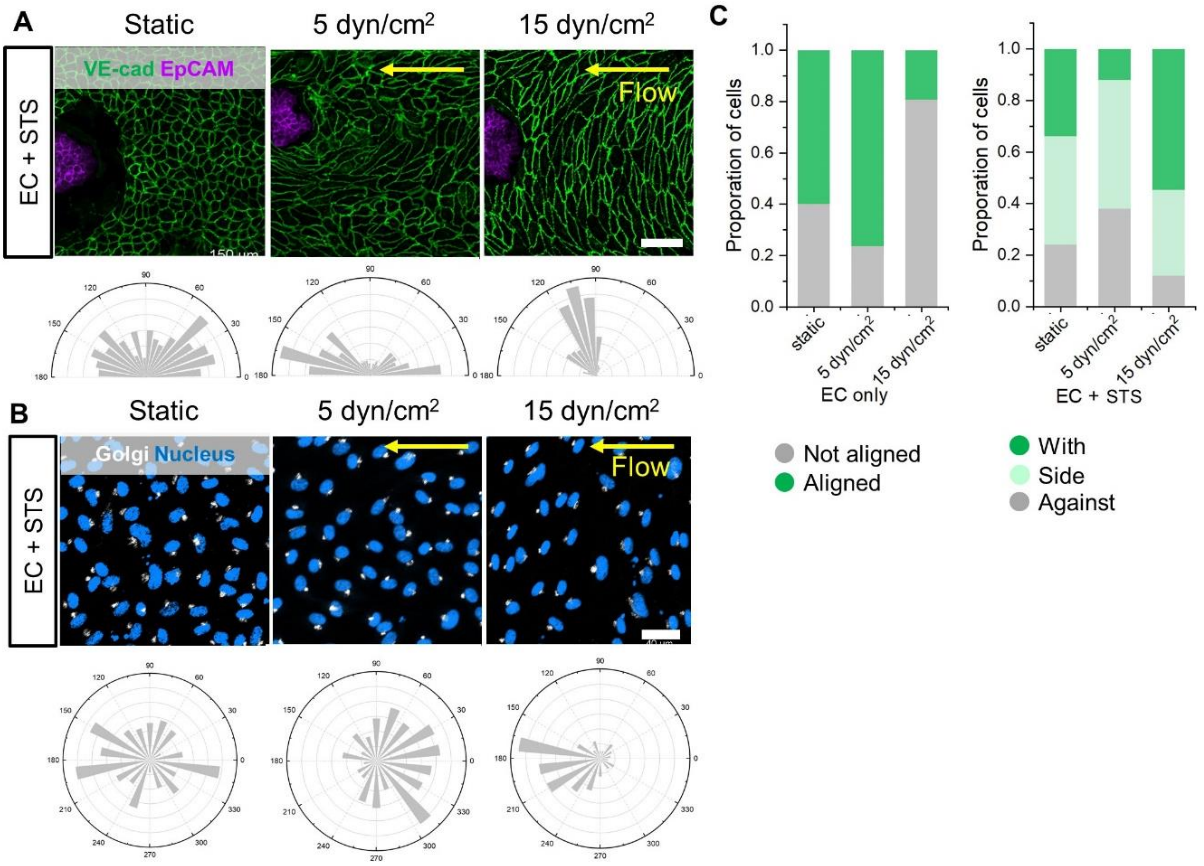


**Supplementary figure 3.** (A) Representative confocal images of ECs with STS exposed to shear stress for 40 hours (static: 0 dyn/cm$^2$; 5 dyn/cm$^2$; 15 dyn/cm$^2$) and associated orientation quantification shown as circular plots. Scale bar = 100µm. (B) Representative confocal images ECs with STS exposed to shear stress for 40 h (static: 0 dyn/cm$^2$; 5 dyn/cm$^2$; 15 dyn/cm$^2$) and associated polarization quantification shown as rose graphs. Scale bars = 40µm. (C) Left: Quantification of the percentage of ECs aligned with the flow direction (in between 45° around the flow axis,) in the presence of STS; Right: Quantification of ECs polarization relative to the flow direction (n = 3; with: in between 135 and 225° around the flow axis, side: 45–135° and 225–315° against 0–45° and 315–360°; approximately 600-700 cells were analyzed).

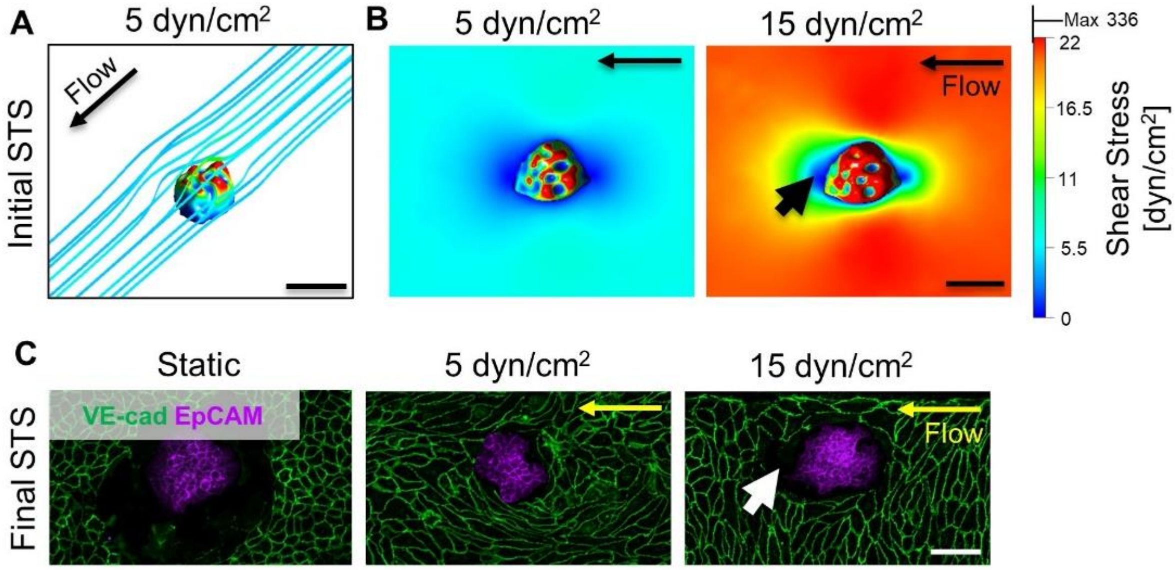


**Supplementary figure 4.** (A) 3D computational fluid dynamic (CFD) simulation of shear rate distribution and flow line around STS. Scale bar = 50µm. (B) The top view of CFD simulation of shear rate distribution of the channel with STS. Scale bar = 50µm. (C) Representative confocal images of STS after 40 hours of static culture (left), exposure to 5 dyn/cm$^2$ (middle) and 15 dyn/cm$^2$ (right). Scale bar = 100µm.

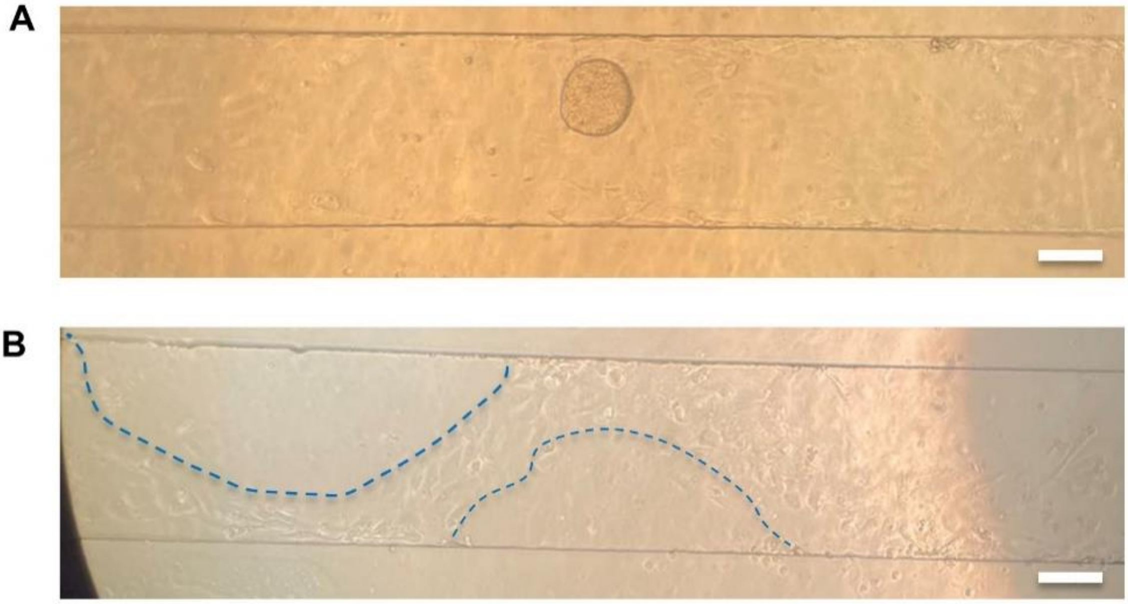


**Supplementary figure 5.** (A) Bright field image of tumor spheroid immediately after being seeded in a microfluidic channel with 300 × 100µm (width × height). Scale bar = 100µm. (B) Bright field image of endothelialized channel (300 × 100µm, width × height) showing that after exposure to shear overnight, the tumour spheroids were detached. The blue dashed lines outline the original positions of the tumor spheroids. Scale bar = 100µm.
